# Fast and flexible simulation and parameter estimation for synthetic biology using bioscrape

**DOI:** 10.1101/121152

**Authors:** Anandh Swaminathan, William Poole, Ayush Pandey, Victoria Hsiao, Richard M. Murray

## Abstract

In systems and synthetic biology, it is common to build chemical reaction network (CRN) models of biochemical circuits and networks. Although automation and other high-throughput techniques have led to an abundance of data enabling data-driven quantitative modeling and parameter estimation, the intense amount of simulation needed for these methods still frequently results in a computational bottleneck. Here we present bioscrape (Bio-circuit Stochastic Single-cell Reaction Analysis and Parameter Estimation) - a Python package for fast and flexible modeling and simulation of highly customizable chemical reaction networks. Specifically, bioscrape supports deterministic and stochastic simulations, which can incorporate delay, cell growth, and cell division. All functionalities - reaction models, simulation algorithms, cell growth models, partioning models, and Bayesian inference - are implemented as interfaces in an easily extensible and modular object-oriented framework. Models can be constructed via Systems Biology Markup Language (SBML) or specified programmatically via a Python API. Simulation run times obtained with the package are comparable to those obtained using C code - this is particularly advantageous for computationally expensive applications such as Bayesian inference or simulation of cell lineages. We first show the package’s simulation capabilities on a variety of example simulations of stochastic gene expression. We then further demonstrate the package by using it to do parameter inference on a model of integrase enzyme-mediated DNA recombination dynamics with experimental data. The bioscrape package is publicly available online (https://github.com/biocircuits/bioscrape) along with more detailed documentation and examples.

## Introduction

In the fields of systems and synthetic biology, it has become increasingly common to build mathematical models of biochemical networks. In principle, such models allow for quantitative predictions of the behavior of complex biological systems and efficient testing of hypotheses regarding how real biological networks function. Such predictions would transform the way in which we design and debug synthetic engineered biological circuits. Many different scales of modeling exist, which are appropriate for different applications. Molecular dynamics models may be used to understand detailed structure-function relationships involving effects at the atomic level [1]. Colloidal simulations coarse-grain individual atoms in order to illuminate the physical principles by which small groups of macromolecules interact [2]. Reaction-diffusion models further reduce the geometry of each molecule to a single point in order to understand spatial and temporal regulation of chemical reactions [3]. Finally, chemical reaction network (CRN) models ignore geometry completely by assuming that the reaction volume is well mixed and focus solely on the chemical reactions that underlie a biological system [4, 5].

CRN models are some of the most widely used in systems, synthetic, and molecular biology. Typically, biological CRN models consist of systems where different species such as DNA, RNA, and proteins can interact with each other via different types of molecular interactions including transcription, translation, activation, repression, and sequestration. Formally, these systems can be represented as a network of species 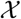 and reactions 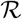. Each reaction 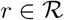 will have the form

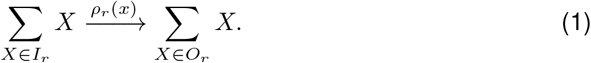

Here *I_r_* and *O_r_* are the input and output multi-sets of species to reaction *r*, respectively. The function *ρ_r_*(*x*) is called a propensity or rate function and controls how quickly the reaction occurs. In general, this can be an arbitrary function of the amount of the species in the network, *x*. Common choices for rate functions include mass action kinetics [5], which are appropriate for detailed mechanistic models, and various kinds of Hill functions for coarse-grained models with sigmoidal saturating reaction rates [5].

A CRN model may be simulated exactly as either a deterministic trajectory of species’ concentrations or a stochastic trajectory of species’ counts. In the deterministic case, the propensity functions can be used to construct a system of ordinary differential equations (ODEs), which can be integrated to understand how a circuit functions in bulk given some initial concentration [6]. This kind of simulation is appropriate when the absolute number of species in the reaction chamber is high, such as in *in vitro* biochemical circuits [7] and in modeling the mean behavior of a population of cells [8]. In the stochastic case, a CRN is simulated as a Markov jump process using the stochastic simulation algorithm (SSA) [9]. Many such simulations can further be combined to understand the time-evolution and steady state of the distribution of the species counts. Biological circuits can often be noisy [10, 11], especially in single cells with low molecular copy numbers [12]. In these cases, a stochastic model is often necessary to capture the noise characteristics of a circuit.

Stochastic simulation also allows for the inclusion of delay into chemical reactions. Processes like protein production are not instantaneous, and there is often a significant delay between when transcription of a gene is initiated and when a mature protein is produced. This type of delay can lead to non-trivial behavior such as oscillations [13], and thus it is often important to incorporate delay into the modeling framework. Additionally, delays can be both fixed and distributed in their duration. While adding a fixed delay to a biological circuit might destabilize the circuit and create oscillatory behavior, distributing that delay across multiple time intervals might maintain circuit stability [14].

Cell growth and division are also critical aspects of biological circuits that operate in single cells. Typically, a dilution term in the model accounts for cell growth. However, in stochastic models, modeling the continuous dilution process with a stochastic and discrete degradation reaction might not be accurate. Another source of noise is the partitioning of molecules between daughter cells at cell division, which can be difficult to distinguish from other forms of noise [15]. Therefore, modeling cell growth as well as division and partitioning is important for investigating noise in gene expression across a lineage of cells.

Regardless of simulation framework, it is necessary to first specify the values of the parameters of each propensity function *ρ_r_*(*x*) in the model along with the initial levels of the model species. In some cases, these parameters and initial conditions are experimentally known. Often, however, they have to be inferred from from biological data via a process known as parameter inference, parameter estimation, or parameter identification [16]. Bayesian inference [17, 18] is one of the most rigorous methods of parameter identification. It provides a posterior distribution over the parameter space so that the stochastic effects from the experimental data are modeled by the parameter distributions instead of a fixed optimal point. This gives insight into the accuracy and identifiability of the model. Also, such an approach allows for an easy comparison between different model classes using the model evidence. The drawback of these approaches is that their implementation is computationally expensive and is based on repeated forward simulations of the model within the framework of Markov chain Monte Carlo (MCMC) [17]. Therefore, it is important to have the underlying simulations running as fast as possible in order to speed up computation time.

Once a given model is fully specified, it is then important to validate the model against additional biological data. In this workflow, it is often necessary to add or remove reactions from the model or to perform a different type of simulation. For example, one might decide that a circuit behaves too noisily for deterministic simulations and want to switch to a stochastic simulation framework. If delays are playing a significant role in the dynamics, one might want to incorporate previously unmodeled delays into the model.

The result is that a very large amount of data is needed to first parameterize and then validate models. The use of technologies for lab automation makes this data collection increasingly accessible and economical. For deterministic models, this may include data collected at many different operating conditions which can be achieved with high throughput measurement techniques involving liquid handling automation [19]. For stochastic models this may include large sample sizes of single cell cell measurements such as flow cytometry [20, 21] and tracking single cell lineages with fluorescent microscopy [22].

This paper presents bioscrape (Bio-circuit Stochastic Single-cell Reaction Analysis and Parameter Estimation), which is a Python package for fast and flexible modeling and simulation of biological circuits. The bioscrape package uses Cython [23], an extension for Python that compiles code using a C compiler to vastly increase speed. This helps assuage the computational time issues that arise in parameter estimation and stochastic simulation. Bioscrape provides an object oriented framework which allows for easily customizable models that can be simulated in many different ways including deterministically, stochastically, or as growing and dividing lineages of single cells. Flexible easy-to-use wrapper and a Python API make it straightforward for a researcher to change their model and try simulations under diverse conditions. Some popular software packages that do somewhat similar tasks as the bioscrape package are MATLAB’s SimBiology toolbox [24] and Stochpy [25]. However, the bioscrape package is faster, supports fully general propensity functions, and allows more kinds of simulation than these alternatives making it more flexible and more efficient than alternative packages.

This paper first details the flexible model specification and simulation capabilities as well as the speed of bioscrape, prior to delving into an example of using bioscrape to perform parameter estimation for integrase enzyme mediated DNA recombination dynamics using experimental data. More detailed documentation and the code for the examples as well the package itself are available online (https://github.com/biocircuits/bioscrape).

## Materials and Methods

### A flexible modeling language for biological circuits

In bioscrape, models can be defined by building the model using Python code, or by importing models in the standardized XML-based language called Systems Biology Markup Language (SBML) [26]. The Python API can be used to model species, reactions (including propensities and delays), and parameter values. Figure 1 illustrates a simple transcription translation model of gene expression, the corresponding Python API model construction, and code necessary to simulate the model with bioscrape. A list of the delay and propensity types supported by bioscrape can be found in Supplemental Table 1 with complete documentation on the bioscrape github Wiki.

**Fig 1.**
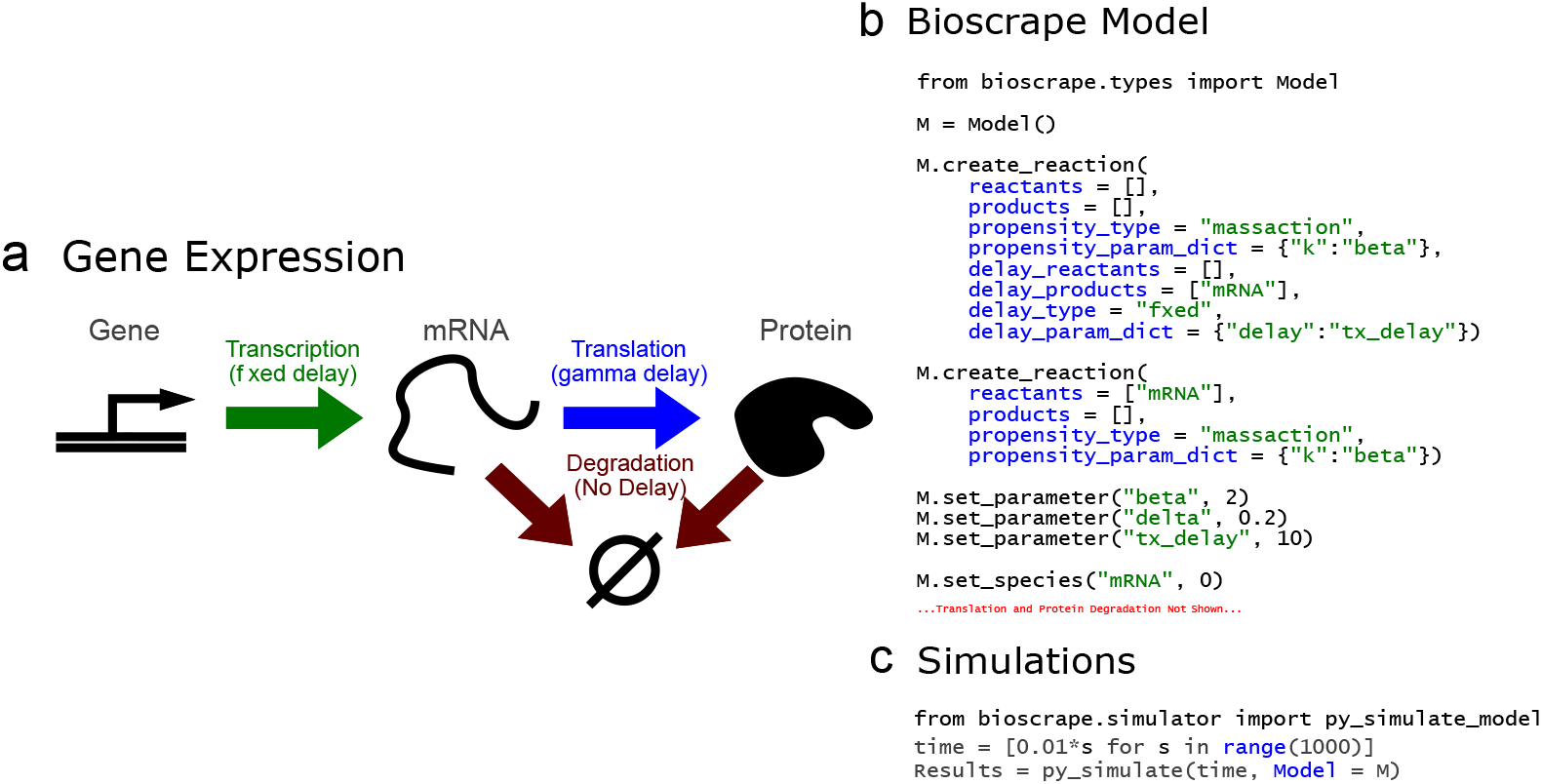
(a) A simple model of gene expression with transcription, translation, mRNA degradation, and protein degradation. The quantity of the gene encoding for mRNA is considered constant and absorbed into the transcription rate *β.* (b) Example Python code to construct a CRN model of gene expression using Bioscrape. (c) Models constructed via SBML or the Python API can be easily simulated with results returned as a pandas Dataframe [27].

### Fast simulation of biological circuits

In addition to the flexible model specification described previously, another critical aspect of a software package for quantitative analysis of biological circuits is speed. This package is written using Cython [23], a language extension for Python that creates compiled Python libraries. Some alternative methods for doing stochastic simulation are to use the SimBiology toolbox in MATLAB [24], write code in C from scratch, or to use a pure Python library such as StochPy [25]. In this section, the simulation speed of the bioscrape package is benchmarked against these other three common simulation options.

The benchmark test used for comparing the speed of these different simulators is a simple gene expression model consisting of just four stochastic reactions: transcription, translation, and degradation of mRNA and protein. The full model is available with the bioscrape package documentation. As MATLAB SimBiology does not support delayed reactions, the system was simulated ignoring delays for 100,000 minutes of simulation time starting from an initial condition of zero. Additionally, both SimBiology and StochPy output each step of the stochastic simulation as opposed to outputting the system state at specific times. For the simulation conditions in this system, the number of steps taken in 100,000 minutes is always around ten million steps. Therefore, to make the comparison fair, the simulation in the bioscrape package is done with ten million desired time points in order to keep the output size similar in all cases. Finally, the C code is a pure C implementation of the simulation using the same fixed interval algorithm as the bioscrape Python package, so the C implementation is also run with ten million desired time points.

The simulation times are available in Table 1. The table shows that bioscrape outperforms SimBiology by almost one order of magnitude, but it outperforms the pure Python StochPy package by a factor of 270. The C simulation is used as a surrogate for the maximum speed possible. The bioscrape package is about twice as slow as custom pure C code. This is due to polymorphism in the way propensities and delays are handled, which greatly improves code readability and flexibility but does cost some speed.

**Table 1.**
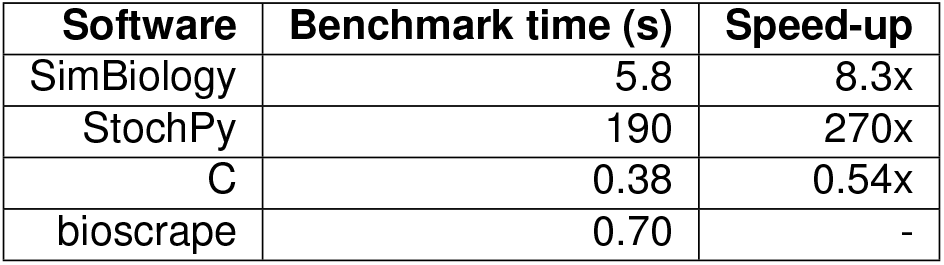
A speed comparison between this Python package and other common simulation platforms.

### Flexible simulation of biological circuits

In addition to speed and flexible modeling, bioscrape also enables flexible simulation of biological circuits. Simulations can be performed in a deterministic or stochastic setting, and stochastic simulations can optionally account for delay as well as cell growth and division. A list of available simulators can be found in Supplemental Table 1 with complete documentation on the bioscrape github Wiki.

Using bioscrape, a simple model of gene expression, illustrated in Figure 1a, is simulated under a variety of different conditions and the results are displayed in Figure 2. This model only contains transcription, translation, and degradation of mRNA and protein as its reactions. In a deterministic simulation, the mRNA and protein levels smoothly trend to their steady state values, while in the stochastic simulations the trajectories are noisy (Figure 2a, b). The delays in the model are a fixed ten minute delay for transcription and a gamma-distributed delay with a mean of ten minutes for translation. When the simulation is performed with delay, mRNA levels spike sharply after a ten minute delay, while protein levels gradually increase at the twenty minute mark. Using the stochastic mRNA trajectories to compute the mRNA distribution as well as the mRNA autocorrelation function results in Figure 2e and Figure 2f, showing that the empirically computed mRNA distribution and autocorrelation match their theoretical counterparts.

**Fig 2.**
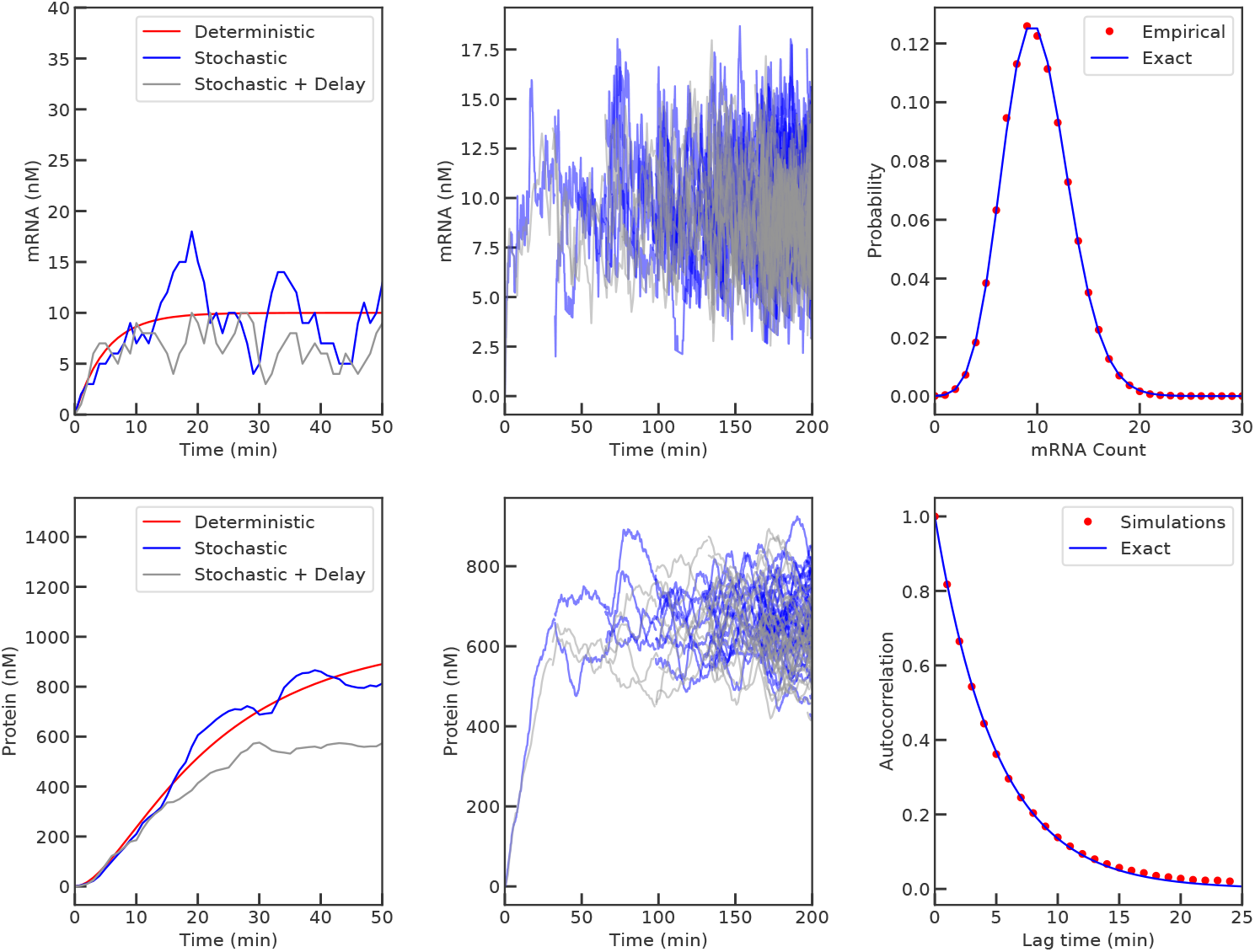
Flexible simulation of a simple model of gene expression with delay. The figure shows mRNA and protein trajectories for the model using a deterministic simulator, a stochastic simulator, and a stochastic simulator accounting for delays. With transcription and translation delays of ten minutes each, mRNA production beings at ten minutes, with protein production starting at twenty minutes. In the second column, the mRNA and protein trajectories across a simulated lineage of cells with and without delay are shown. Steady state mRNA and protein levels are lower with delays in transcription and translation. Finally, the empirical distribution function for mRNA in the simple model of gene expression matches the theoretical Poisson distribution and the empirical autocorrelation function for mRNA in the stochastic simulation matches the theoretical exponential curve.

Because living cells can grow and divide, bioscrape can also simulate biological circuits within the context of a growing and dividing cell lineage (Figure 2c, d). When simulated without delay, the mRNA level in the lineage trends to a steady state of 10 just like in the deterministic simulation, while the protein level trends to its deterministic steady state value of 1000. When delay is introduced, the steady state values of mRNA and protein both decrease. This is because the effective current mRNA or protein production rate with delay is proportional to the number of cells that existed at some time in the past, which in a growing lineage is fewer than the current number of cells. This decreases the effective mRNA and protein production rate per cell, thus decreasing the steady state concentrations.

All of these types of simulations can be performed with a single bioscrape model. Furthermore, toggling between different types of simulations only requires a few lines of code. For example, switching between standard deterministic and stochastic simulations only requires a single line of code, while including volume and cell division requires a few lines of code from the user to specify parameters for cell growth and partitioning of species between daughter cells. All functionalities - simulation algorithms, cell growth models, and partioning models - are implemented as interfaces in an object-oriented manner. This not only ensures ease of switching simulation methods or partitioning models, but also it allows for easy extension of the source code itself to implement new functionality in a modular fashion.

A more in-depth demonstration of the stochastic capabilities of bioscrape, in which bioscrape is used to model the replication and partitioning of plasmids within a growing and dividing lineage of cells, is given in S2 Appendix.

## Results

In the previous sections, we demonstrated the capabilities of bioscrape for performing fast, flexible, and efficient simulations of biological circuits. In this section, we use the package’s parameter inference capabilities to identify the parameter values for a model of enzyme-mediated DNA recombination dynamics based on *in vitro* experimental data. Bioscrape provides a user friendly wrapper to run Bayesian inference on biological system models using a Markov Chain Monte Carlo (MCMC) sampling algorithm [28]. We first start by giving background on integrase systems and *in vitro* prototyping of biological circuits. We then describe the experimental procedure and the experimental data collected. Finally, we introduce the model and perform parameter inference on the model for both simulated data as well as the actual experimental data.

### Background and experimental design

Both serine integrase systems and *in vitro* prototyping using cell free extracts are well-studied tools in synthetic biology. Serine integrases are proteins that can recognize and recombine two specific target DNA sequences [29, 30]. Depending on the original directionality of the target sites, the recombination causes the segment of DNA between the target sites to either be excised or reversed. Figure 3b depicts the process by which four serine integrase monomers bind to attB and attP DNA recognition sites and recombine them into attL and attR sites. In synthetic biology, this functionality has been leveraged to build synthetic gene circuits for state machines [31], temporal event detection [32], and rewritable memory [33]. However, existing applications of integrases rely on their digital behavior over long time scales (hours), and not much is known about the dynamics of their action upon DNA.

**Fig 3.**
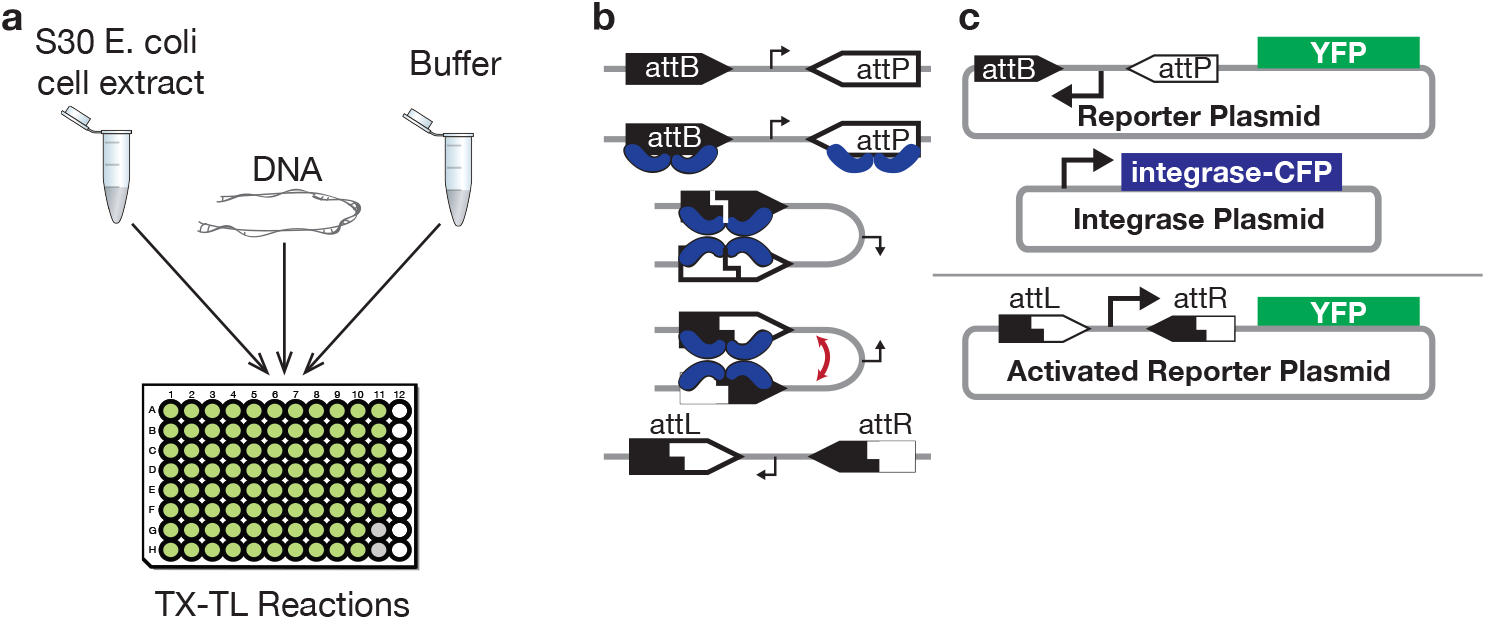
Testing serine integrase recombination dynamics using TX-TL. (a) The TX-TL system allows for prototyping synthetic circuits *in vitro* by adding DNA to cell extract and buffer. (b) Four serine integrase monomers cooperate to recombine attB and attP DNA sites to form attL and attR sites while reversing the DNA segment between the sites. (c) A constitutive integrase expression plasmid expresses integrase fused to cyan fluorescent protein (CFP), which flips a promoter on a a reporter plasmid and leads to yellow fluorescent protein (YFP) expression.

One way to assay the dynamics of integrase DNA recombination is to test an integrase system using TX-TL, an *E. coli* cell extract *in vitro* system for testing and protoyping synthetic gene circuits [34]. Plasmid or linear DNA encoding the genes in a synthetic circuit can be added to a TX-TL master mix to prototype genetic circuits outside the cell as depicted in Figure 3a. In this case, we can create a simple synthetic circuit involving constitutive integrase production and reporter expression following DNA recombination to assay DNA recombination as a function of integrase levels. The circuit consists of two plasmids as shown in Figure 3c. On the first plasmid, the integrase plasmid, we constitutively express Bxb1, a commonly used serine integrase, as a part of a fusion protein in which Bxb1 is fused to CFP (cyan fluorescent protein). This allows us to use CFP fluorescence to measure the amount of Bxb1 present in the TX-TL reaction. The second plasmid is a reporter plasmid in which a promoter initially pointing away from a yellow fluorescent protein (YFP) gene can be reversed by integrase DNA recombination to point towards the YFP gene, which leads to production of YFP. Therefore, YFP expression can be used to infer when DNA recombination has occurred. Detailed plasmid maps are available in S4 Appendix.

### Experimental Results

Using automated acoustic liquid handling (Labcyte Echo 525), we varied the level of integrase plasmid and reporter plasmid between 0 and 1 nM across 100 different TX-TL reactions. Each reaction contained integrase and reporter plasmid both independently at one of five concentrations of 0 nM, 0.25 nM, 0.50 nM, 0.75 nM, or 1 nM. Therefore, there were 25 possible combinations of concentrations of the two plasmids. Four replicates were done for each combination of concentrations, yielding a total of 100 TX-TL reactions. The reactions were incubated at 37 degrees Celsius, and CFP and YFP fluorescence were collected every 5 minutes for each reaction using a plate reader. Using a previously performed calibration of fluorescence to concentration, we were able to convert the fluorescence measurements for CFP and YFP to actual concentrations in nM. Notably, the CFP concentration allowed us to measure the concentration of Bxb1 integrase in the reaction.

Figure 4a shows the median expression over time for integrase and reporter for each combination of concentrations of integrase and reporter plasmid. Each column of plots corresponds to a reporter plasmid concentration, while darker lines correspond to increasing integrase plasmid concentration. As expected, increasing integrase plasmid increases CFP expression. With no reporter or no integrase plasmid, YFP expression is absent as expected. It is also clear that reporter expression generally begins sooner and ends at a higher level when there is more integrase expression. The full set of experimental data is available in S1 Appendix.

**Fig 4.**
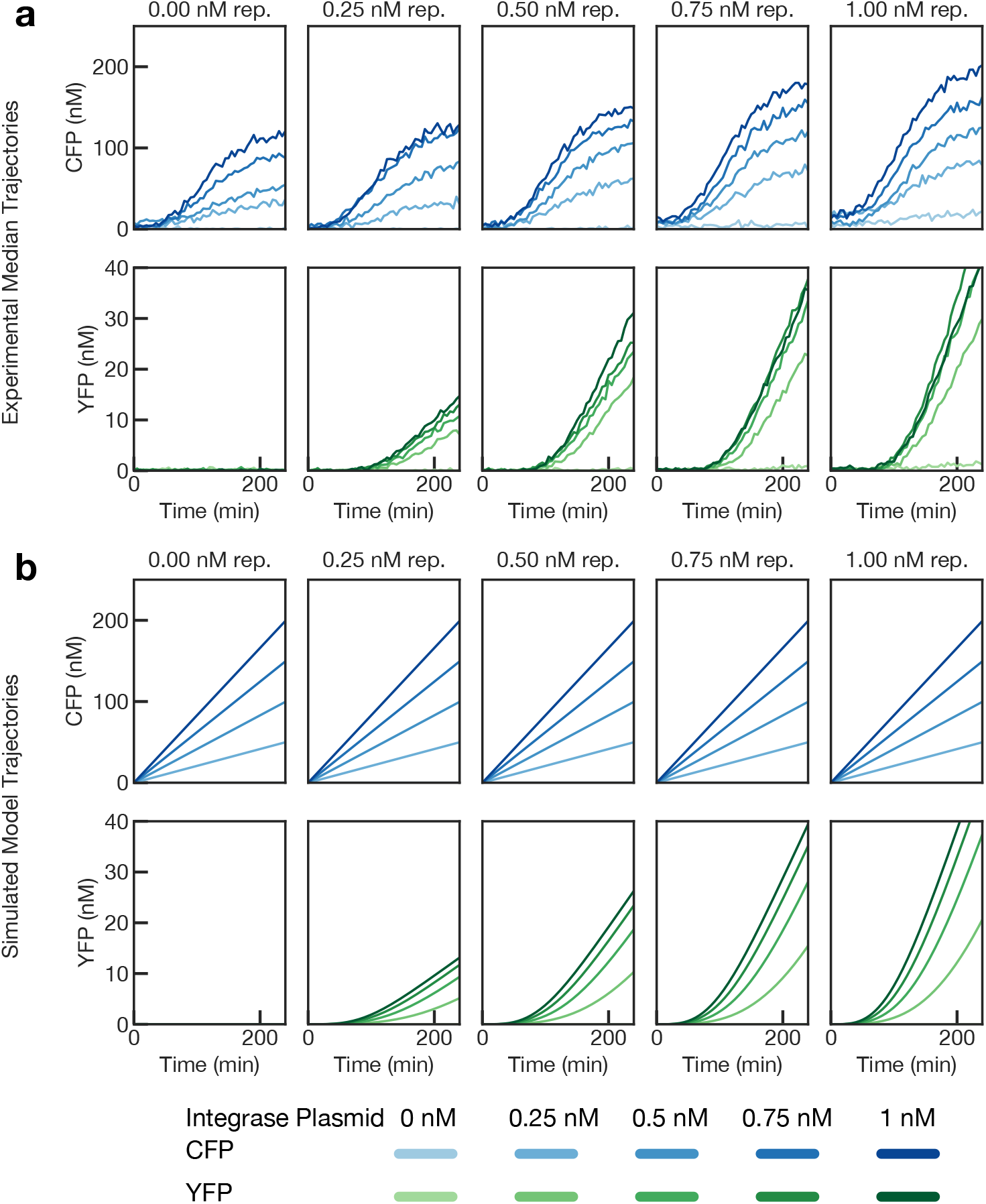
Experimental and model simulation results. (a) Both integrase plasmid and reporter plasmid were varied from 0 to 1 nM and fluorescence data was collected for 4 hours. The plotted lines are the median of 4 replicates per condition. Each pair of vertical plots represents a concentration of the reporter plasmid. Darker lines correspond to higher concentrations of the integrase plasmid. CFP expression corresponds to integrase concentration, while YFP corresponds to reporter. (b) A simulated version of panel a using the model.

### Model of integrase recombination

In order to estimate parameters for the integrase data presented in the previous section, we need a model of integrase recombination of DNA. We created a simple phenomenological model of integrase dynamics consisting of three reactions: integrase production, DNA recombination, and reporter production. As TX-TL is a bulk environment, we use a deterministic model for our system, which we easily set up using bioscrape.

In Table 2, we describe the species in the model. These species are then used in the following set of ODE’s that describe the integrase recombination dynamics in the model. The ODE model may be obtained by first writing a chemical reaction network description [35] then using a formal model reduction procedure [36] to obtain the Hill function in the model description:

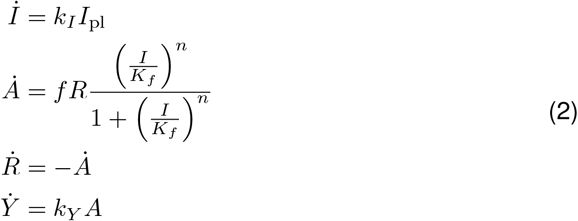

**Table 2.** Species and parameters in the simple model of integrase recombination

| Variable | Species |
| --- | --- |
| $I$ | Integrase-CFP (nM) |
| $A$ | Activated reporter plasmid (nM) |
| $R$ | Unactivated reporter plasmid (nM) |
| $Y$ | YFP fluorescent reporter (nM) |
| $I_{pl}$ | Integrase plasmid (nM) |
| Parameter | Description |
| $k_I$ | Rate of integrase production (nM integrase per minute per nM integrase plasmid) |
| $f$ | Maximum rate of integrase flipping of DNA (nM activated plasmid per nM reporter plasmid per minute) |
| $K_f$ | Hill threshold for integrase activation (nM integrase) |
| $n$ | Hill coefficient |
| $k_Y$ | Rate of reporter production (nM reporter per minute per nM activated plasmid) |

Equation 2 contains the ODE’s for the simple model. Integrase is produced at a constitutive rate, where *I*_pl_ is the concentration of integrase plasmid and varies across experiments. The conversion of reporter plasmid to activated reporter plasmid is governed by a Hill function that allows us to model the cooperativity and activation threshold for the integrases in a simple way. We also assume that the DNA recombination reaction is first order in reporter plasmid. Finally, we assume that YFP reporter is produced at a rate proportional to the amount of activated reporter plasmid. While varying the integrase plasmid changes the value of *I*_pl_ in the model, varying the reporter plasmid changes the initial condition for *R.*

Using representative values for the model, we created a simulated version of Figure 4a using the model. The plot is given in Figure 4b, and there are some qualitative differences between integrase expression in the simulations and in the experimental data. Namely, while in the model the expression of integrase increases linearly with a slope proportional to the amount of integrase plasmid, in the experimental data, integrase expression only increases after a delay and then levels off after about two hours. This behavior is common in cell free extracts due to depletion of resources, and this effect should be included in a future more detailed model of the system. The full model for integrase dynamics is included with the bioscrape package documentation.

### Parameter inference for integrase dynamics

Using the model given in Equation 2, we attempted to perform parameter inference using bioscrape to fit the model parameters to both the simulated data from Figure 4b as well as the experimental median data from Figure 4a. Fitting the model to simulated data was an empirical test of the identifiability of the model from the collected data. If a simulated version of the data were uninformative about parameter values in the models, then the real data would not be informative about the parameters either.

Bioscrape provides a easy-to-user wrapper for parameter inference wherein the user simply needs to import the model and the experimental data as a CSV file. Bayesian inference algorithm in bioscrape is implemented using an off-the-shelf ensemble Markov chain Monte Carlo package that generally works well on parameter inference problems [28, 37]. With bioscrape, various tunable settings for Bayesian inference can be easily changed, such as, the number of steps of the MCMC chain, the number of walkers, the parameters to identify, and the initial seeds for the parameters. Further, bioscrape provides built-in likelihood computation objects for different kinds of biological data and prior distributions. The user may simply specify the prior distribution for each parameter by choosing from one of the built in probability distributions or define their own custom prior function. A customized prior probability distribution can play an important role in an iterative scheme of parameter inference where the posteriors from one run are used as priors for the next run. A detailed list of all priors available with bioscrape are available on the Wiki page alongside detailed documentation of the same. Demonstrative examples for common parameter inference scenarios such as a linear fit, a birth-death model, experimental replicates, varying initial conditions, and delayed models are also available with the bioscrape package.

Figure 5 contains the posterior distributions for the parameters obtained after performing parameter estimation on the simulated data. It is clear that the posterior parameter distributions are tightly centered around the maximum aposteriori estimate values, demonstrating that the parameters are identifiable from the simulated data. This can be considered an empirical identifiability analysis that is necessary for parameter estimation on the experimental data to be meaningful.

**Fig 5.**
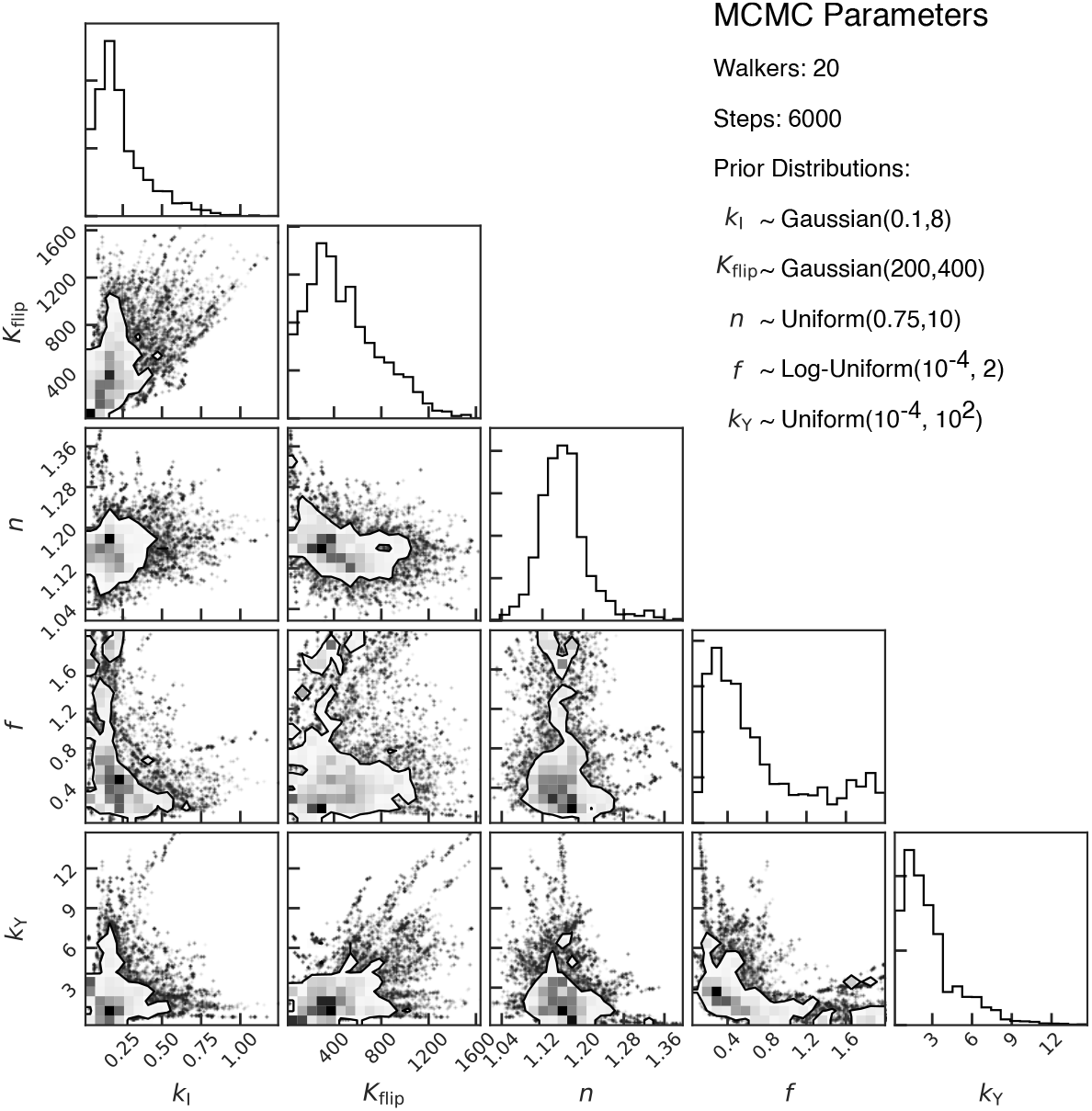
Posterior parameter estimates from simulated data using MCMC. Parameter distributions are shown along with their covariance. The inference using simulated data gives us an empirical insight into the identifiability of the model.

Figure 6 contains the posterior parameter estimates from parameter estimation on the experimental data. In this case, the production rate parameters *k_I_*, *f*, and *k_Y_* have strongly peaked posterior distributions. However, the Hill coefficient *n* and the and the Hill activation constant *K_f_* have a broader range across the space. Moreover, the posterior samples for the activation constant *K_f_* have peaked at higher values compared to the simulated data. This suggests that integrase recombination is quite slow in practice. Details on the parameter estimation procedure and the comparison of simulated trajectories generated by the estimated parameters to the experimental data are available in S3 Appendix.

**Fig 6.**
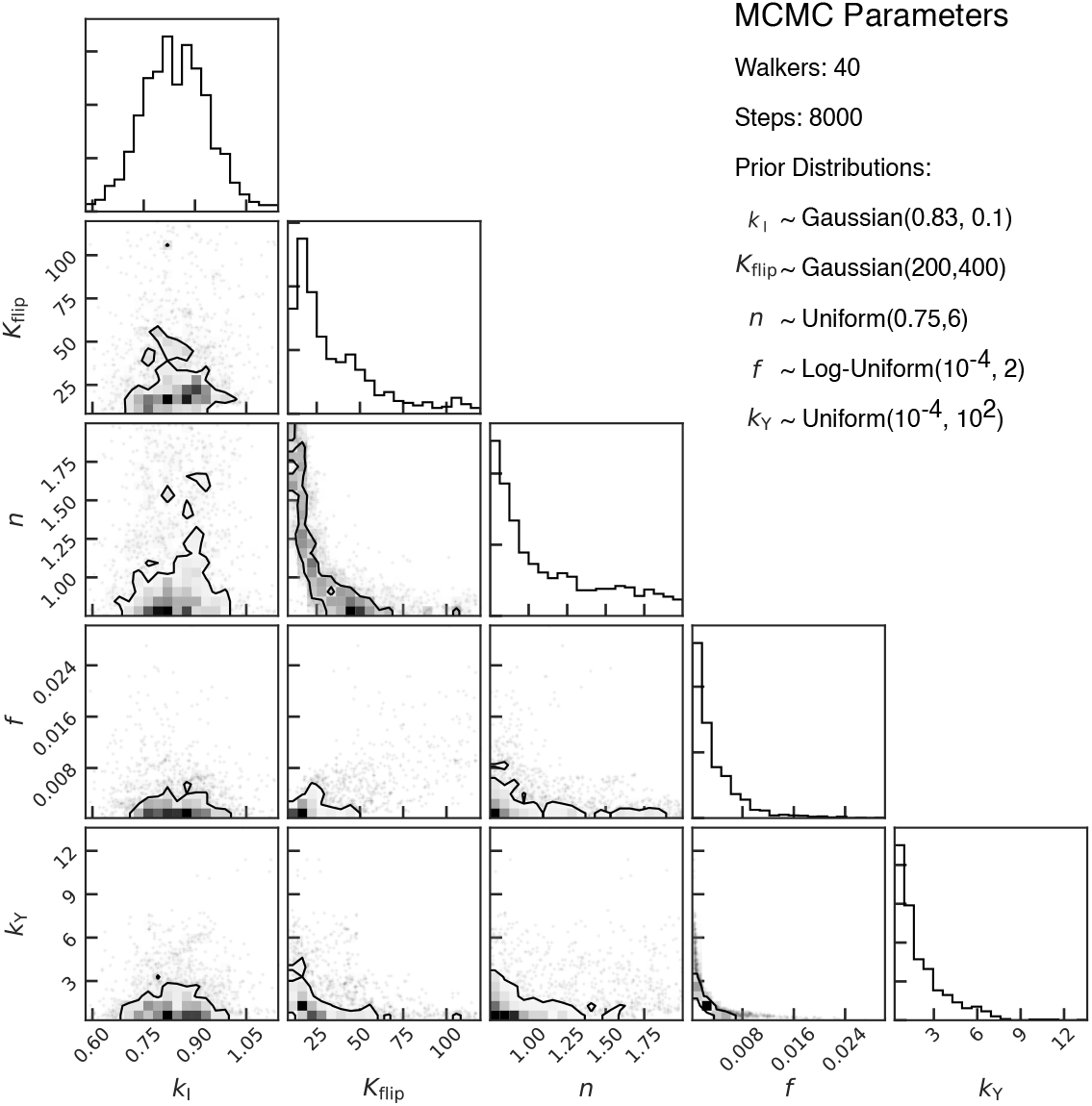
Posterior parameter estimates from experimental data using MCMC. Parameter distributions are shown along with their covariance. The covariance plots show the probability level sets for each pair of parameters where the parameter values inside the contour lie within 60% of the maximum aposteriori estimate. The prior distributions were chosen in an iterative manner by using posterior of MCMC run with initial uninformative priors to inform the stricter prior distributions for the next run. The total run time for each iteration of Bayesian inference with bioscrape was 24 minutes on a machine with Intel i9 CPU and 32 GB of RAM.

In general, the amount of data required for identifiability and parameter estimation to work will vary on a case-by-case basis. Our recommendation is that the user always perform a simulated version of the experiment and parameter estimation on the simulation results to determine whether parameter estimation is feasible in each scenario. However, even if parameter estimation works well on the simulated data, it may still not work well on the experimental data. This may occur because the model does not perfectly describe reality, or because there is noise in the system and measurements. These errors can particularly confound estimation of the less sensitive parameters in the model, such as the Hill coefficient above.

## Availability and Future Directions

The advent of increased computational resources and high-throughput data collection for biological circuits has made quantitative modeling and parameter estimation for biological circuits more feasible. Since the most attractive parameter estimation techniques rely on Bayesian inference and Markov chain Monte Carlo (MCMC), it is important to have a simulator that can perform fast forward simulations of the model. Additionally, the simulator must be able to produce the same types of data that are observed in standard biological assays such as flow cytometry or fluorescence microscopy, so that simulated and experimental data can be compared. Also, as models often need to be tweaked to fit the data, it should be easy to change the model or the way the model is simulated (e.g. switching from a deterministic to a stochastic simulation).

The bioscrape package addresses all of these issues. The flexible model specification using Python allows a user to easily make modifications to a biological circuit model. A library of simulators is available for performing simulations and easily swapping between deterministic and stochastic simulations as well as consideration of other common effects in biological circuits such as cell growth, division, and delays. Finally, because the package is written in Cython, its speed is comparable to the speed obtained using C code.

Performing simulations that incorporate effects such as cell growth, division, and delay can provide insight into the behavior of biological circuits. For example, in this paper, it is demonstrated that in an exponentially dividing colony of cells, delays in gene expression can lead to a lower steady-state protein concentration.

However, the ultimate aim of bioscrape is to provide tools for doing parameter estimation for synthetic and systems biology. Here, we demonstrated the use of the bioscrape package to perform parameter estimation for both simulated and experimental data for integrase recombination dynamics in the TX-TL cell-free *in vitro* system. As a result of this demonstration, we were able to estimate dynamic parameters for integrases that may be relevant for synthetic circuit design.

The fast simulators presented here will be the computational workhorse for more complex MCMC schemes, enabling parameter inference on stochastic models of synthetic gene circuits. Future work and future updates to the software will include inference methods and an experimental demonstration for a stochastic model.

## Acknowledgments

AS, AP, and VH were supported by the Defense Advanced Research Projects Agency (Agreement HR0011-17-2-0008). The content of the information does not necessarily reflect the position or the policy of the Government, and no official endorsement should be inferred. AS was also supported by AFOSR grant FA9550-14-1-0060. AP was also supported by the NSF grant CBET-1903477. WP was supported by an NSF Graduate Research Fellowship (No.2017246618).

The authors acknowledge members of the Murray lab at Caltech for assistance with experiments and helpful feedback and also acknowledge all the members of the scientific community at large who have used and provided feedback on bioscrape.

## Supporting information

**Supplemental Table 1:**
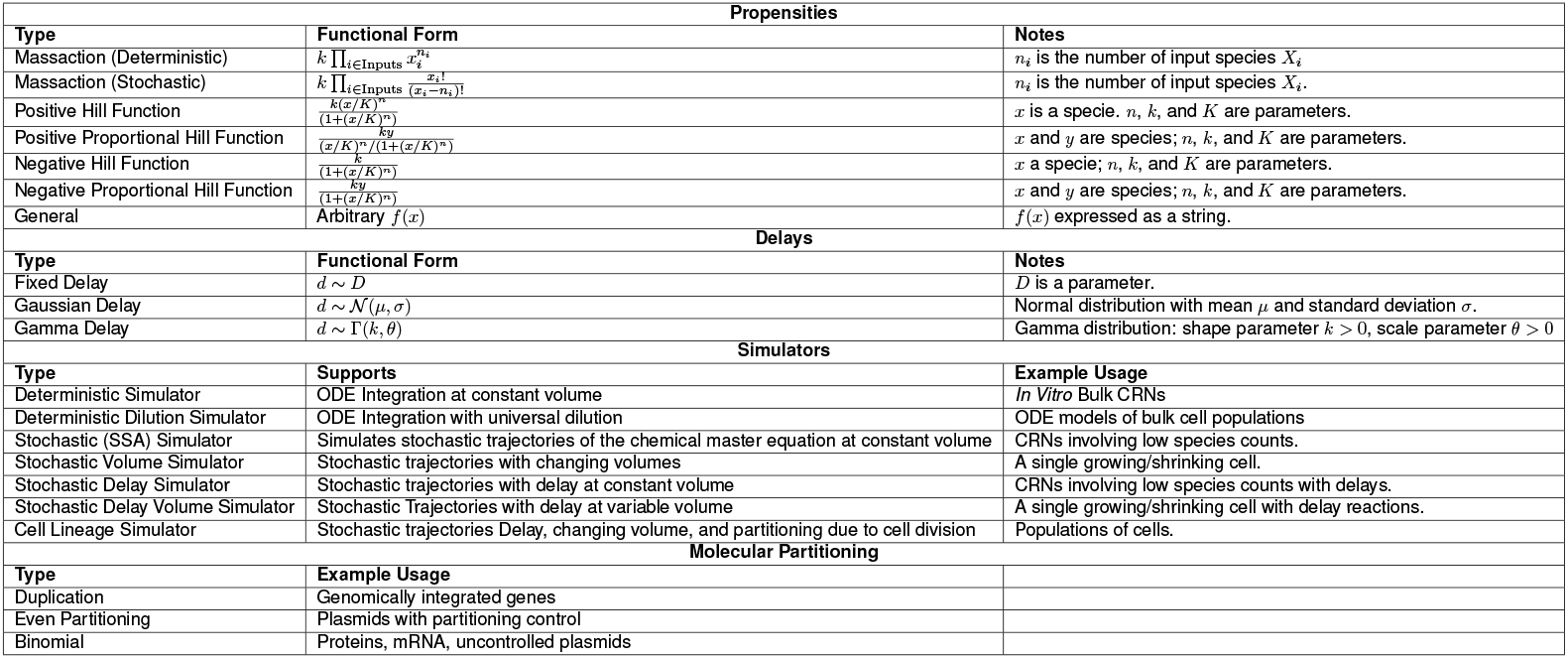
Bioscrape Features

## S1 Appendix. Complete TX-TL data

**Fig 7.**
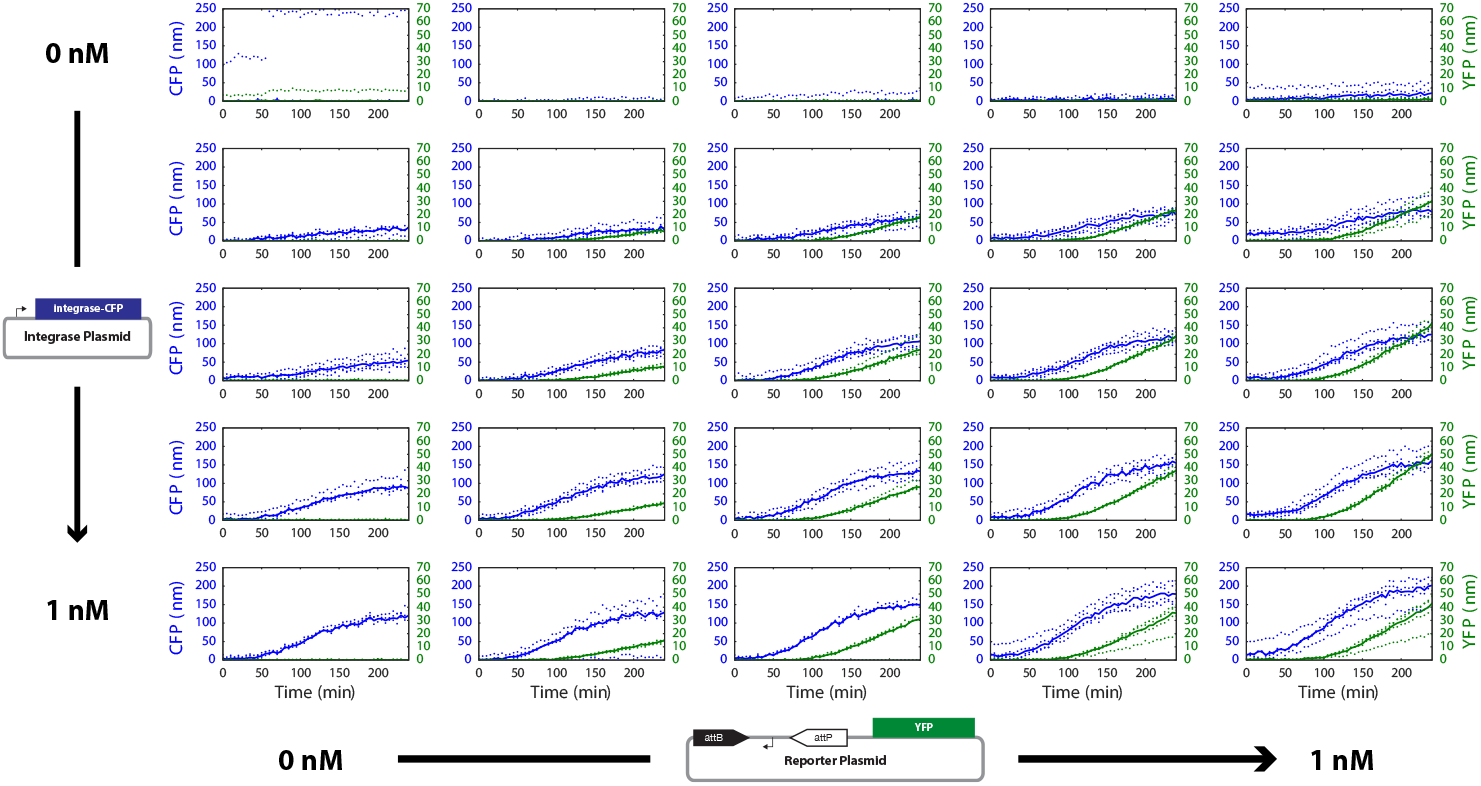
The complete TX-TL data used to generate the median expression plots in Figure 4a. Each plot corresponds to a concentration of integrase plasmid and reporter plasmid. Dots correspond to measured data points, and solid lines correspond to the median of four replicates.

## S2 Appendix. A reduced order model for plasmid replication in single cells

Using a simplified model of plasmid copy number control that is derived here, bioscrape is used to model a system of plasmids that replicate and partition randomly between daughter cells but also transcribe an mRNA. In doing so, bioscrape simulation results can be used to model noise in transcription accounting for noise in plasmid copy number in addition to intrinsic transcriptional noise.

The ColE1 plasmid regulates its own copy number by constitutively transcribing an RNA that inhibits the RNA primer for DNA replication from initiating a replication event [38]. Making a four simplifying assumptions enables the derivation of a simplified model of plasmid copy number regulation. First, it is assumed that the inhibitory RNA directly binds to the plasmid origin to inhibit replication. Second, it is assumed that that the replication rate is proportional to number of free plasmids, which do not have inhibitory RNA bound. Third, it assumed that the inhibitory RNA transcription and degradation dynamics are much faster than the plasmid replication dynamics. Fourth, the inhibitory RNA is assumed to be strongly transcribed and linearly degraded, so that the steady state level of inhibitory RNA is much greater than the number of plasmids. The third and fourth assumptions enable the inhibitory RNA to be considered as being at a quasi-steady state level.

Given *P* copies of plasmid, the third and fourth assumptions above mean allow for the steady state level of inhibitory RNA *R* to be approximated by *kP*, where *k* is a large proportionality constant.

Then, assuming fast binding and unbinding of the RNA to and from the plasmid with some dissociation constant *K_d_,* the following equations describing dissociation and mass conservation must hold.

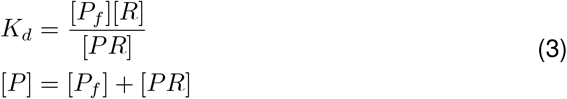

Here, [*P*] denotes the concentration of *P*, so 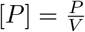, where *V* is the cell volume. The variable *P_f_* denotes the number of free plasmids, while *PR* is the number of plasmid-RNA complexes, which have to add up to the total number of plasmids. Solving these two equations yields the following expression for [*P_f_*].

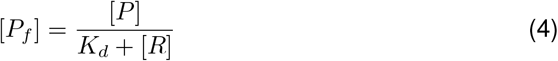

Here, since *k* ≫ 1, [*R*] will be mostly unaffected by its binding to the plasmid, so substituting the steady state expression of *R* gives the following expression for [*P_f_*].

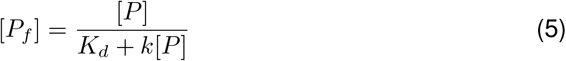

The initiation rate of plasmid replication is assumed to be proportional to the amount of free plasmids *P_f_*, so multiplying both sides by the volume and re-arranging variables gives

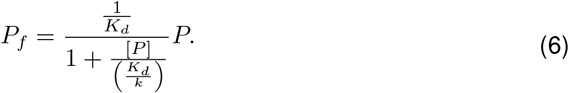

Since the propensity of plasmid replication is assumed to be proportional to the number of free plasmids *P_f_*, the variables can be re-arranged to write down the following expression for the replication propensity, where the parameters have been combined into two parameters *β* and *K*.

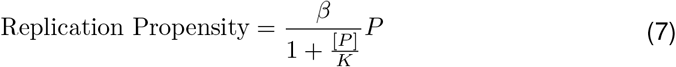

A deterministic analysis of this plasmid replication rate can be performed. To do this analysis, assume that the cell volume is growing at a standard exponential rate with

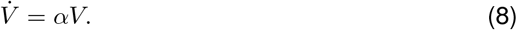

Then, the dynamics of [*P*] can be computed.

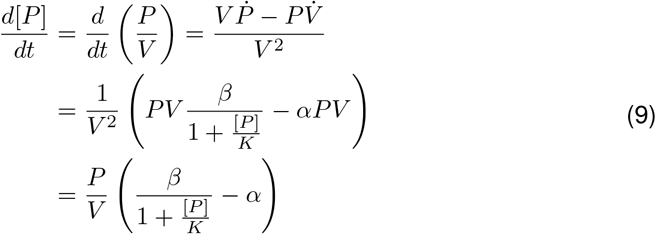

Setting the derivative equal to zero and solving gives the steady state value for the plasmid concentration.

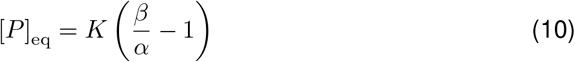

If volume is measured in units of cellular volume, then the average plasmid concentration can be thought as the steady state plasmid copy number. Note that *β* > *α* is required in order to have a non-negative steady state plasmid concentration. This is because the maximum rate of plasmid production must at least be able to keep up with the cell growth rate in order for the plasmid to be maintained.

### Simulating plasmid replication and gene expression in single cells

Using the model of plasmid replication derived in the previous section, a model of plasmid replication combined with transcription can be used to compute the variability in mRNA levels between cells in a lineage simulation. In the model, there is one plasmid species, which replicates itself and also constitutively transcribes a mRNA. It is possible to look at the plasmid copy number and mRNA levels in a cell lineage over time as well as the plasmid copy number distribution across a population of cells at the end of the simulation. The full model used for producing the simulation is available with the bioscrape package documentation. However, the model is tuned to produce a mean plasmid concentration of 10 nM, and the cell division time is set to 33 minutes, which is a representative division time for *E. coli.*

The simulation is performed for 500 minutes and the plasmid distribution is empirically calculated using a final population size of 2048 cells. The run time for this simulation to compute a total of 4095 cell traces is less than two seconds on a standard desktop computer without using parallel processing.

As shown in Figure 9, the copy number at the end of the simulation has a wide distribution with a mean of about 15 copies per cell. This is expected because the mean concentration should be about 10 nM for the plasmid and the mean cell volume will be around 1.5 volume units. There is a slight peak in the distribution at a copy number of zero. This is because if a cell loses all its plasmids, it will continue dividing but its future descendants will never be able to recover the plasmid.

**Fig 8.**
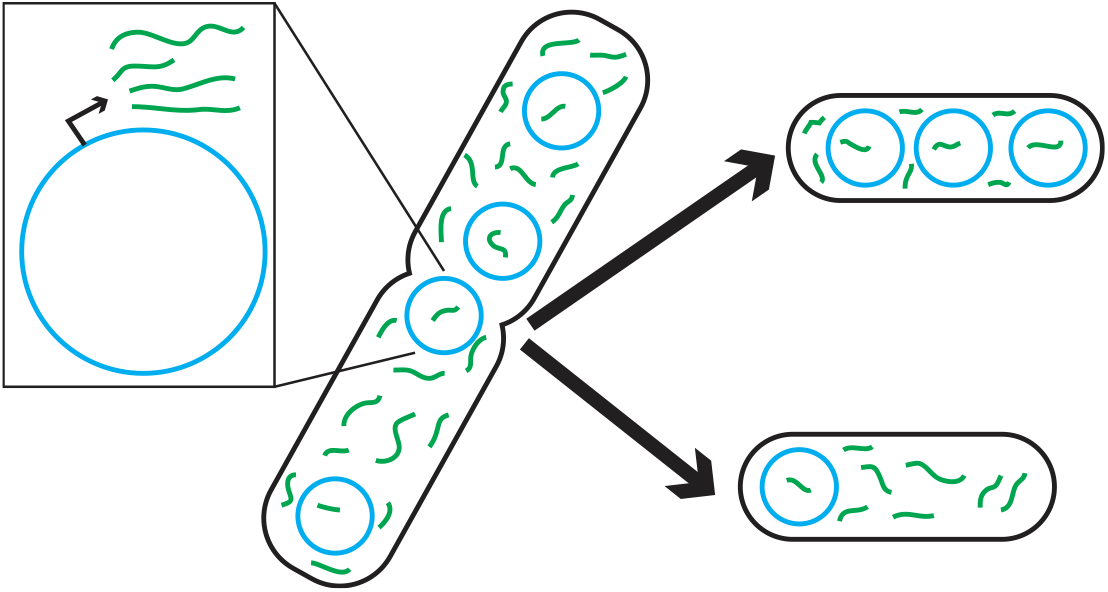
Plasmid transcription with partitioning. Each plasmid (blue) constitutively transcribes green RNA molecules. Both plasmids and RNA’s are partitioned between daughter cells during cell division.

**Fig 9.**
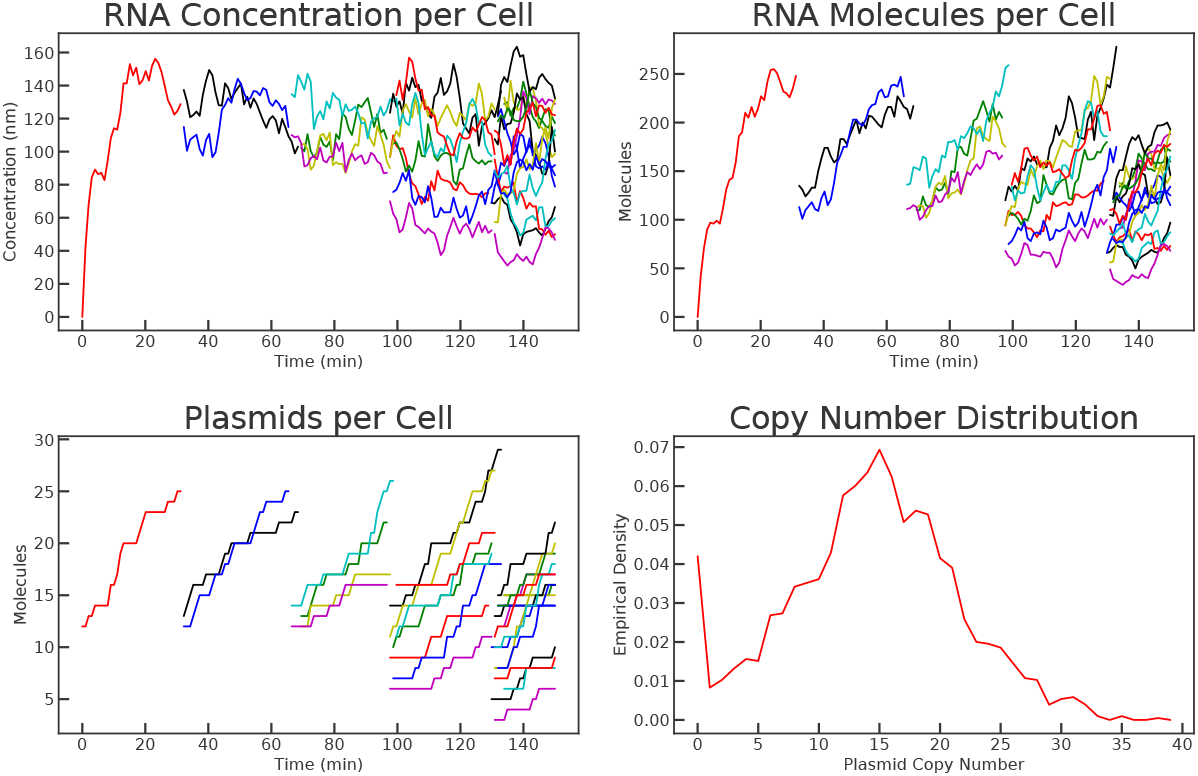
A simulation of plasmid replication and transcription over a cell lineage. The first three plots show trajectories of RNA and plasmid counts and concentrations over time. The last plot shows the distribution of plasmid copy number over 2048 cells at the end of the simulation.

The distribution of plasmid and mRNA concentrations can also be plotted. In this case, the copy number is divided by the cell volume at the end of the simulation before plotting. The expected plasmid concentration is 10 nM and the expected mRNA concentration is 93.45 nM. The results can be seen in Figure 10.

**Fig 10.**
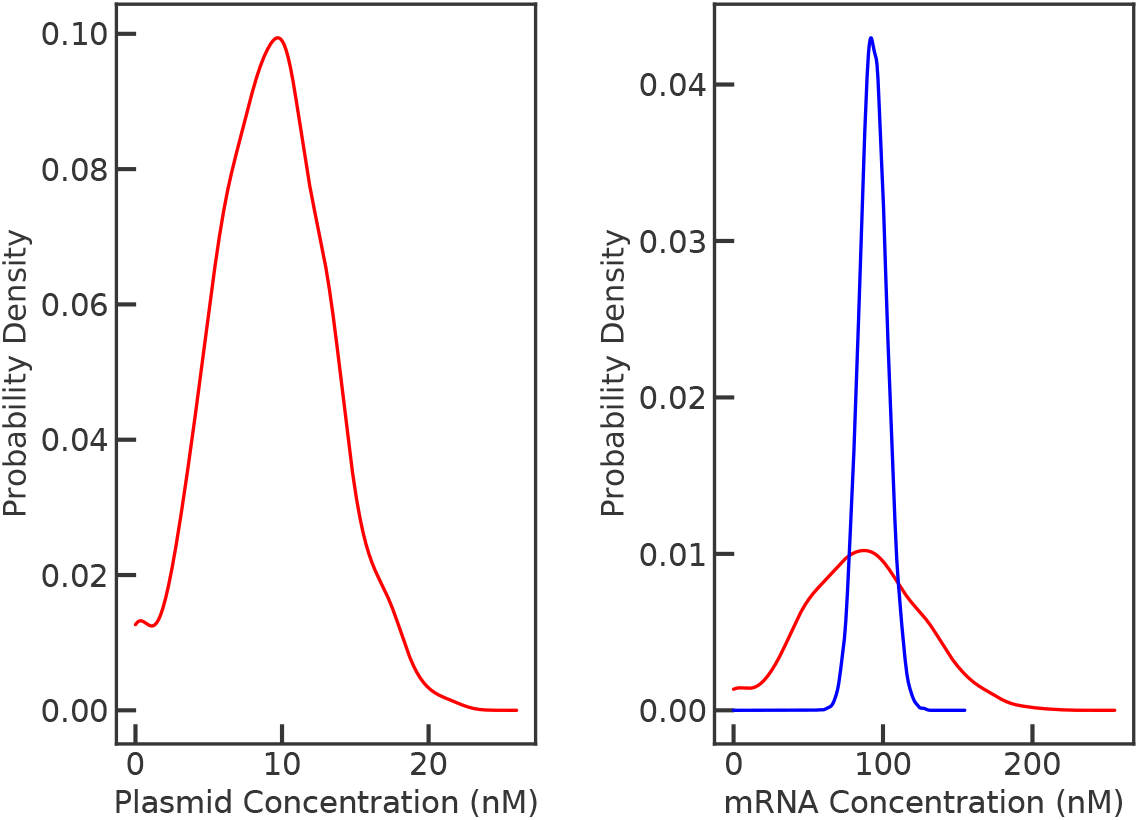
Distributions of plasmid and mRNA concentration across a cell lineage. The plasmid concentration is distributed around a mean of 10 nM. mRNA concentration is distributed around a mean of approximately 90 nM. The blue line shows the mRNA expression distribution if the plasmid concentration was exactly its mean value of 10 nM at all times.

The right panel of Figure 10 also shows a control where the plasmid concentration is assumed to be exactly controlled within the cell with no variability. In this case, the noise in mRNA expression is much smaller than in the case where the mRNA is expressed from the plasmid. The coefficient of variation in the plasmid based expression case is 0.55, while the coefficient of variation in the case with controlled copy number is 0.10.

## S3 Appendix. Details of parameter estimation and model fit

We performed parameter estimation using an ensemble method [37]. Using either the simulated model data or the experimental data as input, we used a least squares log-likelihood function, in conjunction with a different prior distributions over a reasonable space of parameter values. The parameter inference process was completed over multiple iterations to explore the parameter space effectively. For the first few iterations, we initialized the MCMC method with 40 walkers assigned to random positions within a valid but uninformative parameter space. We ran this chain for 8000 iterations. With these iterations, we were able to create prior distributions for each parameter using the posterior of the previous iteration. Each such iteration for the parameter inference using the experimental data took around 24 minutes on an i9 3.10 GHz CPU with 32 GB of RAM. Finally, to confirm convergence, we ran a much longer and extensive MCMC run with 400 walkers and 8000 steps as well. This run took a total of around 3 hours to complete.

To get a representative sample from the posterior that matches the observed data the best, we sampled from the obtained posterior distributions as per the recommendation of [37]. This ensemble was used to generate Figure 6. We show the full model fit to median experimental data for CFP and YFP across all experimental conditions in Figure 11 and Figure 12 for a representative set of posterior parameter values.

**Fig 11.**
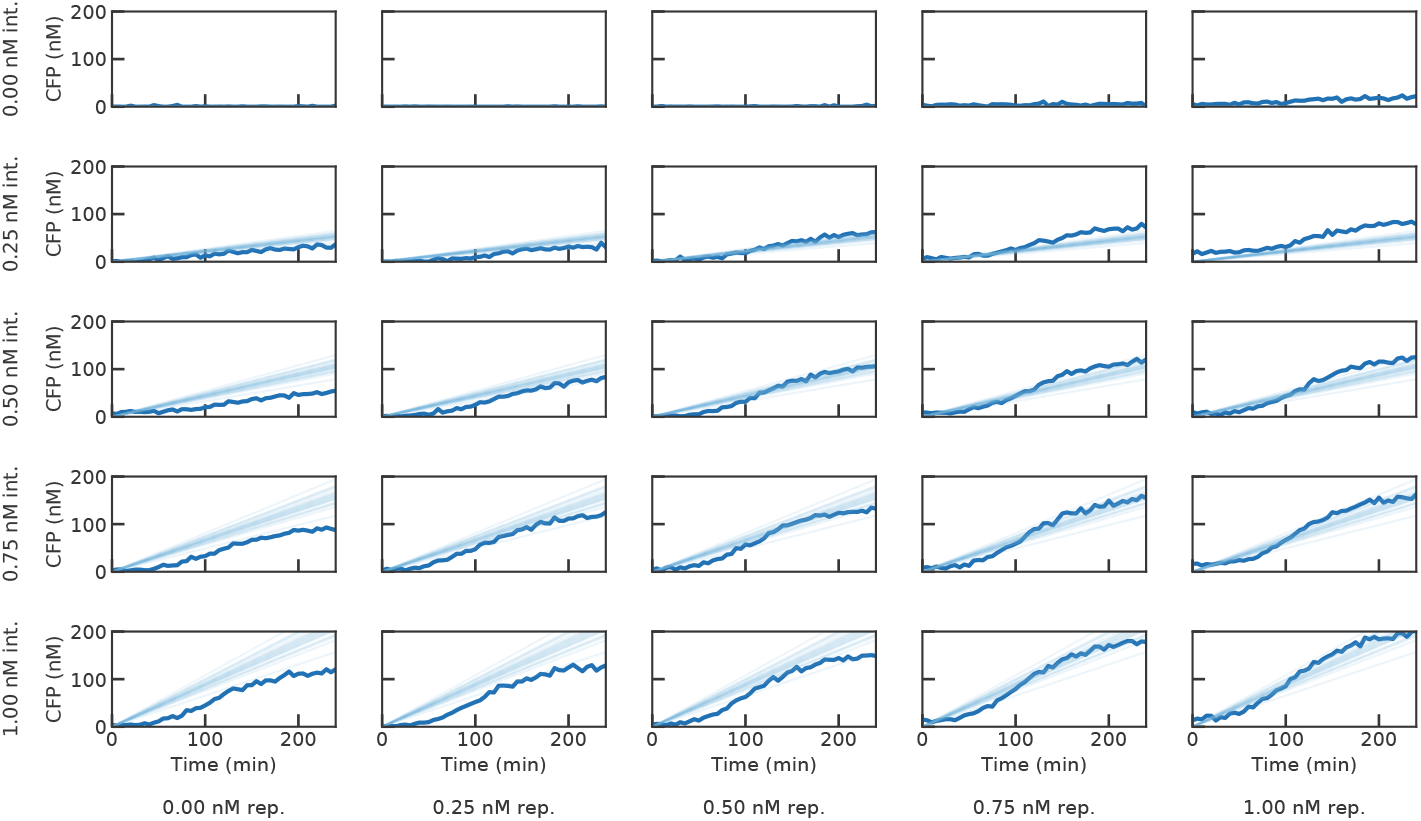
Simulated trajectories using the model with parameters from the posterior distribution (translucent) overlaid on median experimental data (solid lines) for CFP for each combination of integrase and reporter plasmid concentrations.

**Fig 12.**
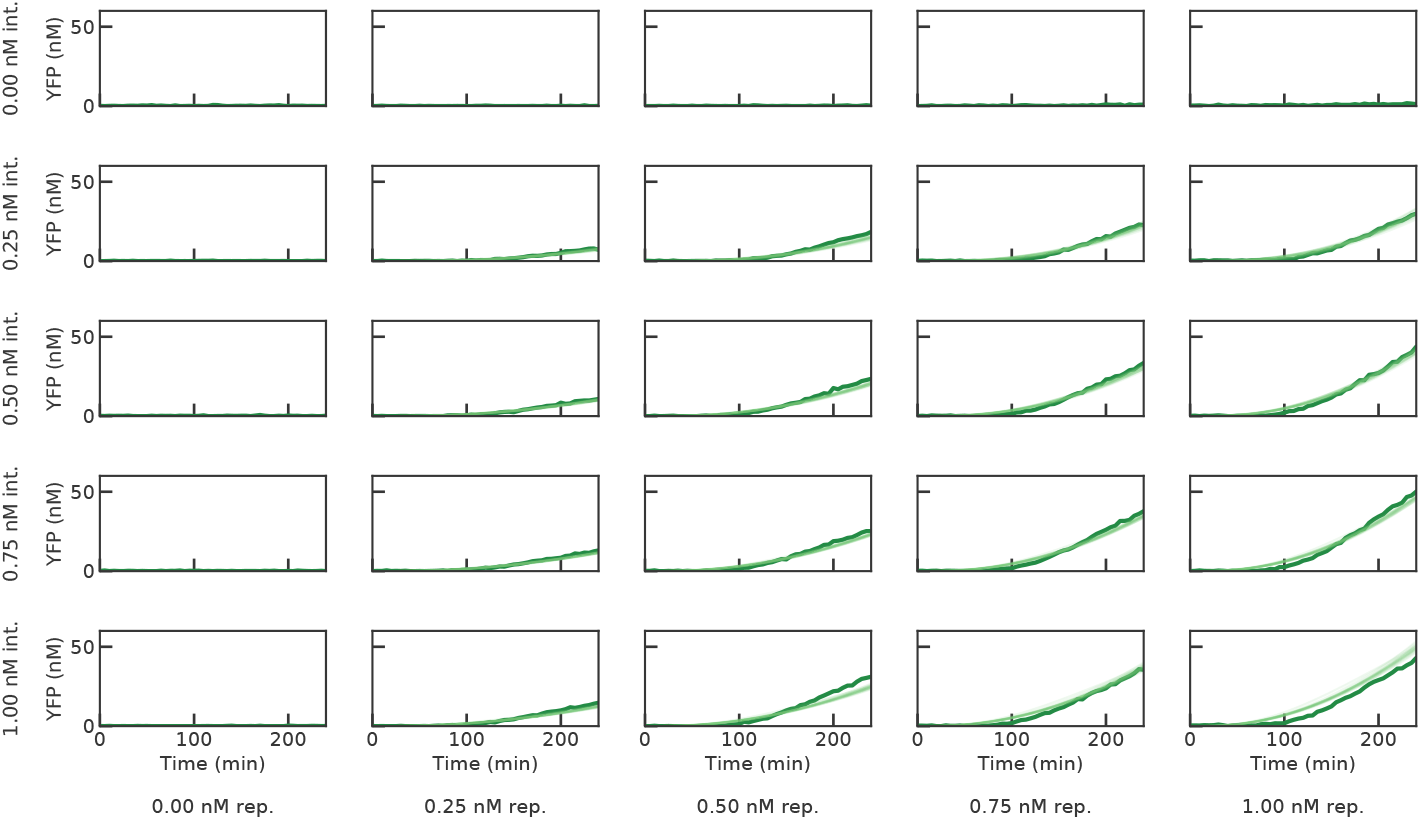
Simulated trajectories using the model with parameters from the posterior distribution (translucent) overlaid on median experimental data (solid lines) for YFP for each combination of integrase and reporter plasmid concentrations.

While YFP trajectories are fit very well by the model, as shown in Figure 12, the experimental CFP trajectories in Figure exhibit some nonlinearity, which cannot be captured by the model. This is likely due to resource limits in cell free extract.

## S4 Appendix. Plasmid maps and experimental details for integrase and reporter plasmids

Experiments in TX-TL were performed and fluorescence data was calibrated to concentration as previously described [39].

The integrase plasmid is illustrated in Figure 13. The plasmid contains a Ptet promoter [40] upstream of a strong ribosome binding site [41], which drives expression of a fusion protein consisting of Bxb1 integrase [33] linked to a cyan fluorescent protein. Transcription is terminated by a T500 terminator [42].

**Fig 13.**
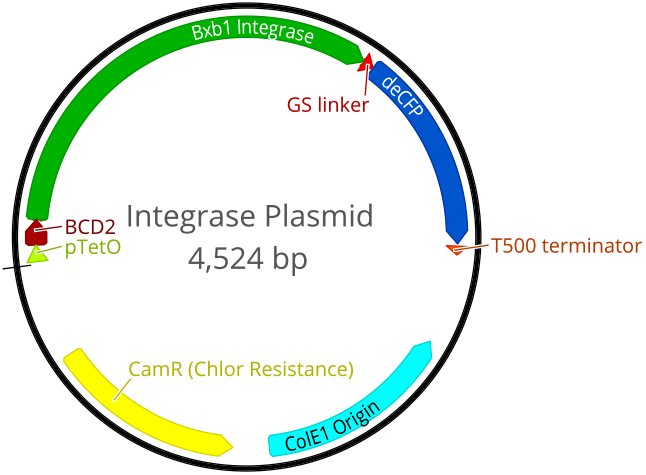
Plasmid map for the integrase plasmid.

The reporter plasmid is illustrated in Figure 14. The plasmid contains a single gene with a strong P7 promoter [41] inbetween Bxb1 recombination sites, so that recombination activates transcription of the gene. A RiboJ insulator [43] and a strong BCD2 ribosome binding site [41] drive expression of the reporter gene, a Venus fluorescent protein.

**Fig 14.**
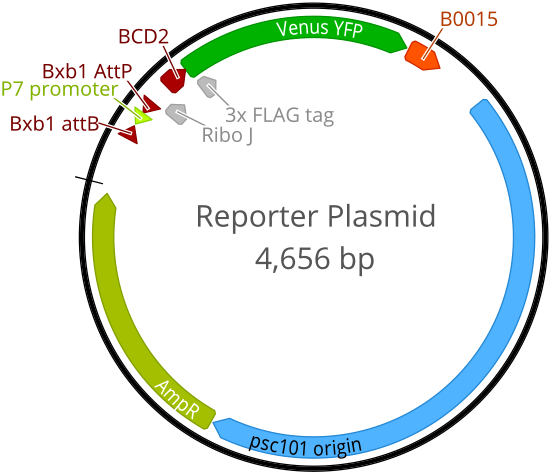
Plasmid map for the reporter plasmid.

